# Research Advance: Extending chemical perturbations of the Ubiquitin fitness landscape in a classroom setting

**DOI:** 10.1101/139352

**Authors:** David Mavor, Kyle A. Barlow, Daniel Asarnow, Yuliya Birman, Derek Britain, Weilin Chen, Evan M. Green, Lillian R. Kenner, Bruk Mensa, Leanna S. Morinishi, Charlotte Nelson, Erin M. Poss, Pooja Suresh, Ruilin Tian, Taylor Arhar, Beatrice Ary, David Bauer, Ian Bergman, Rachel Brunetti, Cynthia Chio, Shizhong Dai, Miles Dickinson, Susanna Elledge, Cole Helsell, Nathan Hendel, Emily Kang, Nadja Kern, Matvei Khoroshkin, Lisa Kirkemo, Greyson Lewis, Kevin Lou, Wesley Marin, Alison Maxwell, Peter McTigue, Douglas Myers-Turnbull, Tamas Nagy, Andrew Natale, Keely Oltion, Sergei Pourmal, Gabriel Reder, Nicholas Rettko, Peter Rohweder, Daniel Schwarz, Sophia Tan, Paul Thomas, Ryan Tibble, Jason Town, Kaitlyn Tsai, Fatima Ugur, Douglas R. Wassarmann, Alexander Wolff, Taia Wu, Derrick Bogdanoff, Jennifer Li, Kurt S. Thorn, Shane O’Conchúir, Danielle L. Swaney, Eric D. Chow, Hiten Madhani, Sy Redding, Daniel N. Bolon, Tanja Kortemme, Joseph L. DeRisi, Martin Kampmann, James S. Fraser

## Abstract

Although the primary protein sequence of ubiquitin (Ub) is extremely stable over evolutionary time, it is highly tolerant to mutation during selection experiments performed in the laboratory. We have proposed that this discrepancy results from the difference between fitness under laboratory culture conditions and the selective pressures in changing environments over evolutionary time scales. Building on our previous work (Mavor et al 2016), we used deep mutational scanning to determine how twelve new chemicals (3-Amino-1,2,4-triazole, 5-fluorocytosine, Amphotericin B, CaCl_2_, Cerulenin, Cobalt Acetate, Menadione, Nickel Chloride, p-fluorophenylalanine, Rapamycin, Tamoxifen, and Tunicamycin) reveal novel mutational sensitivities of ubiquitin residues. We found sensitization of Lys63 in eight new conditions. In total, our experiments have uncovered a sensitizing condition for every position in Ub except Ser57 and Gln62. By determining the Ubiquitin fitness landscape under different chemical constraints, our work helps to resolve the inconsistencies between deep mutational scanning experiments and sequence conservation over evolutionary timescales.

**Builds on:** Mavor D, Barlow KA, Thompson S, Barad BA, Bonny AR, Cario CL, Gaskins G, Liu Z, Deming L, Axen SD, Caceres E, Chen W, Cuesta A, Gate R, Green EM, Hulce KR, Ji W, Kenner LR, Mensa B, Morinishi LS, Moss SM, Mravic M, Muir RK, Niekamp S, Nnadi CI, Palovcak E, Poss EM, Ross TD, Salcedo E, See S, Subramaniam M, Wong AW, Li J, Thorn KS, Conchúir SÓ, Roscoe BP, Chow ED, DeRisi JL, Kortemme T, Bolon DN, Fraser JS. Determination of Ubiquitin Fitness Landscapes Under Different Chemical Stresses in a Classroom Setting. *eLife*. 2016.

**Impact Statement:** We organized a project-based course that used deep mutational scanning in multiple chemical conditions to resolve the inconsistencies between tolerance to mutations in laboratory conditions and sequence conservation over evolutionary timescales.

## Introduction

Ubiquitin (Ub) is an essential eukaryotic protein acting as post-translational modification to mediate the degradation of ~80% of the proteome and regulate protein localization and activity. (Yau and Rape, 2016). The amino acid sequence of ubiquitin has been strikingly stable throughout evolutionary time: between yeast and human, there are only 3 amino acid changes (96% sequence identity) (Finley et al., 2012; Roscoe et al., 2013). However, deep mutational scanning experiments in yeast have revealed that Ub is surprisingly robust to sequence changes, with 19 positions freely mutating to almost any other amino acid without a loss of fitness (Roscoe et al., 2013).

To interrogate the dichotomy between the mutational robustness and the strong sequence conservation of Ub, we initially hypothesized that sensitivities to mutations at new positions could be revealed by growing yeast under different selective pressures in a classroom setting. To test this idea, we previously determined the fitness landscape of ubiquitin in four different chemical perturbations (DTT, caffeine, hydroxyurea (HU), and MG132) (Mavor et al., 2016). We showed that three of the perturbations (DTT, caffeine and HU) sensitize a shared set of positions to mutation. Under standard laboratory culture conditions, these fitness defects are buffered and undetectable. Conversely, we showed that the proteasome inhibitor MG132 increases the mutational robustness of the ubiquitin sequence landscape. Inhibiting the proteasome reduces protein turnover through the same pathway as mutations in ubiquitin, leading to an alleviating interaction between MG132 and many of the mutant alleles. However, 12 of the 19 residues of the tolerant face defined by (Roscoe et al., 2013) of Ub could still be freely mutated, suggesting that the environmental conditions constraining these positions were not sampled by these four perturbations.

To build on these results, we again involved the first-year graduate students in UCSF’s iPQB and CCB programs to determine the fitness landscape of ubiquitin in distinct environments. We chose twelve new chemical perturbations (3-Amino-1,2,4-triazole (3-AT), 5-fluorocytosine (5-FC), Amphotericin B (AmpB), CaCl_2_, Cerulenin, Cobalt Acetate (Cobalt), Menadione, Nickel Chloride (Nickel), p-Fluorophenylalanine (p-FP), Rapamycin, Tamoxifen, and Tunicamycin), which impose a wide range of stresses upon the cell, including osmotic shock, protein folding stress, and DNA damage. Our results represent an important next step towards how deep mutational scanning can be used to explain the evolutionary constraints on sequence conservation patterns.

## Results

### Distinct chemical treatments can sensitize or increase robustness of Ub to mutation

We calculated distribution of fitness changes using the EMPIRC-BC method (Mavor et al. 2016) across each pairwise environmental comparison (**Figure 1**) and examined the response of individual residues (**Figure 2**). Treatment with 3-AT, AmpB, CaCl_2_, Cobalt, and p-FP shifted the residual distribution towards the reduced fitness (left shift in **Figure 1**) when compared to the distribution of two DMSO treatments. The sensitizing compounds (3-AT, AmpB, CaCl_2_, Cobalt, and p-FP) have a dominant effect at positions near hydrophobic patch residues (8, 44, 70) and the C-terminus (**Figure 2**), which is similar to that previously observed after treatment by Caffeine, DTT, or HU (Mavor et al. 2016).

**Figure 1:**
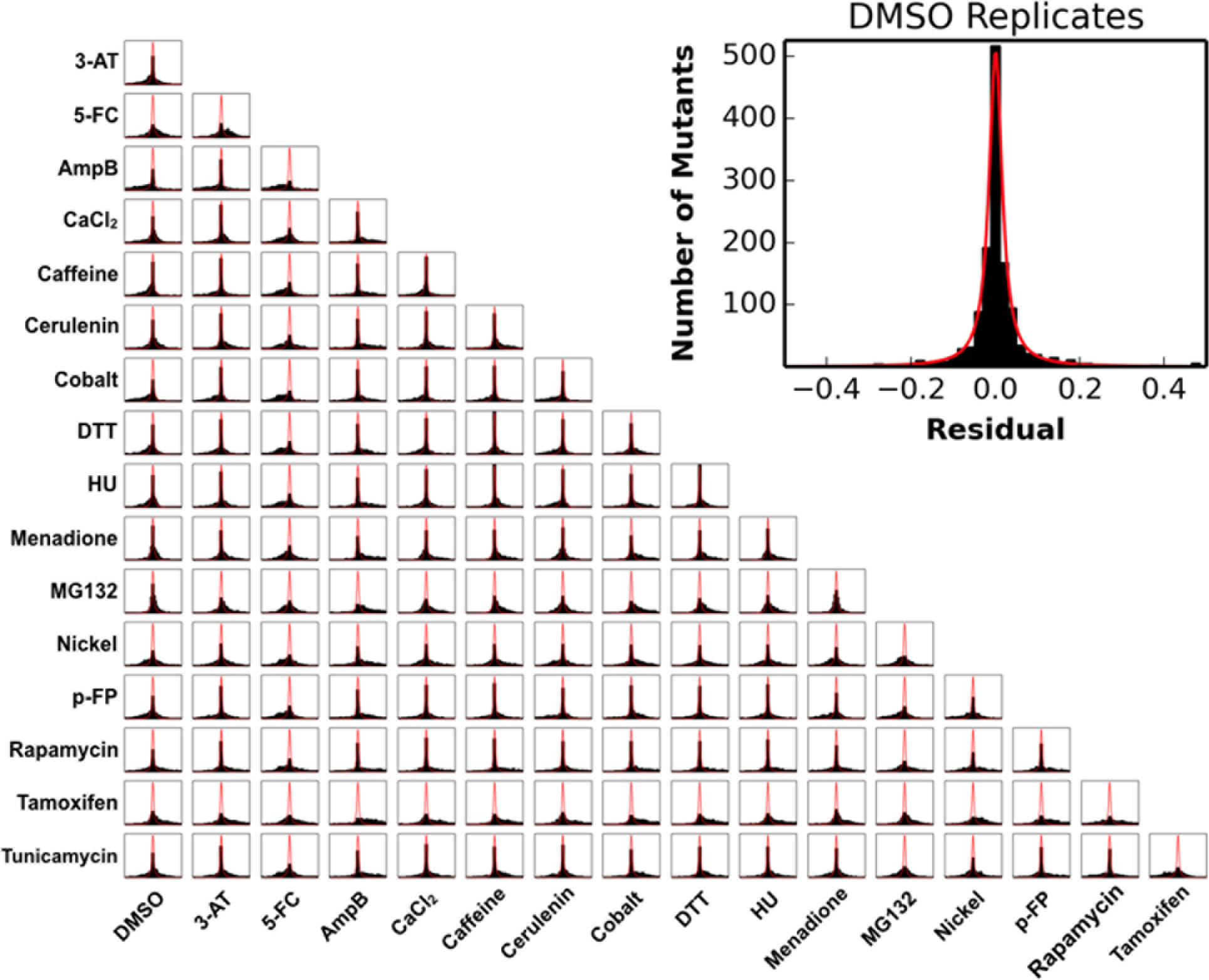
The residuals between data sets reveal global patterns: Inset shows the Lorentzian fit to biological replicates of DMSO in red and data in black. When treatments are compared to DMSO (first column), 3-AT, AmpB, CaCl_2_, Cobalt, and p-FP shift the residual distribution to the left. This highlights the increased sensitivity to mutation. In contrast, 5-FC, Cerulenin, Menadione, and Tamoxifen shift the distribution to the right, highlighting the strong alleviating interactions caused by these treatment. For Nickel, Rapamycin, Tunicamycin we observe long tails on either side of the distribution, highlighting that these treatments induce both strong sensitizing and strong alleviating effects on mutations to Ub.

**Figure 2:**
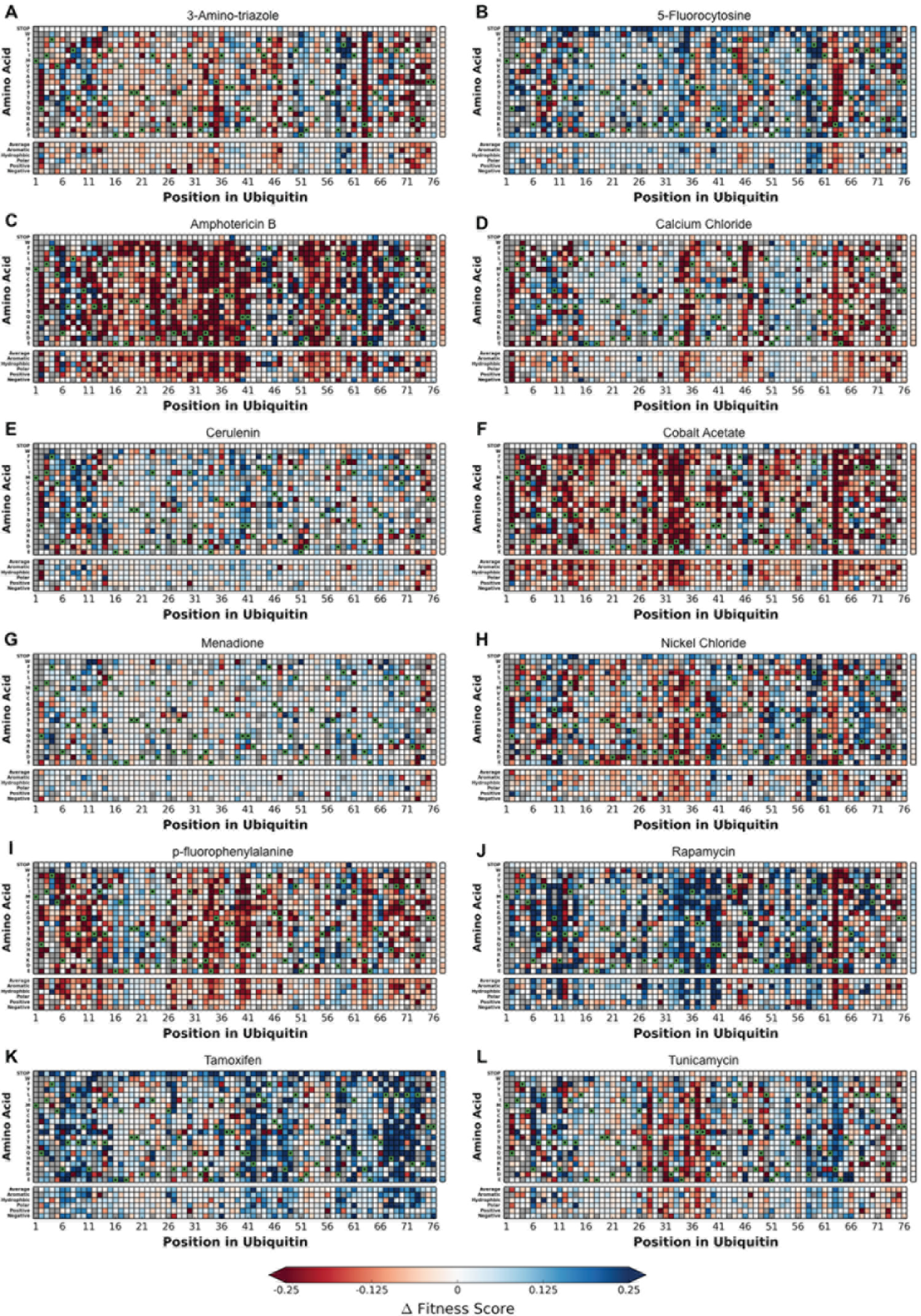
The difference in fitness between DMSO and a perturbation for each Ub allele: (A) 3-AT, (B) 5-FC (C) AmpB (D) CaCl_2_ (E) Cerulenin (F) Cobalt (G) Menadione (H) Nickel (I) p-FP (J) Rapamycin (K) Tamoxifen (L) Tunicamycin. Difference in fitness is represented from 0.25 (Blue) to −0.25 (Red) with white representing no change from DMSO. Wild type amino acids are shown in green and mutations without fitness values (due to lack of barcode or competition sequencing reads) are shown in grey.

In contrast, 5-FC, Cerulenin, Menadione, and Tamoxifen led to more residues having fitness greater than in DMSO (right shift in **Figure 1**). Treatment with Cerulenin or Menadione acted similarly to treatment with MG132; most positions in Ub have mildly increased robustness to mutation, with a few specific mutations appearing either sensitive or tolerant (**Figure 2**). However, 5-FC and Tamoxifen show strong mutational robustness at the biologically important positions (Mavor et al 2016) adjacent to the hydrophobic patch and C-terminus (**Figure 2**).

We observed distinct fitness landscapes for Nickel, Rapamycin, Tunicamycin. These treatments caused large tails in the distribution towards increased and decreased fitness, suggesting that these chemicals induce a unique pattern of sensitization and robustness for different mutants (**Figure 1**). For example, positions 40 and 58 become more tolerant to mutation when cells are exposed to Nickel, but positions 32 through 35 become strongly sensitized. Additionally, several positions show a diverse response to mutation to specific residue types when cells are treated with Nickel (**Figure 2**). For example, Gln2 has an increased relative fitness when mutated to Asn or a positive residue, but a decreased relative fitness when mutated to small hydrophobic or negative residues. Rapamycin and Tunicamycin have many residues where the mutational effects mirror each other (**Figure 2**). For positions 2, 32, 35, 37, and 38 Rapamycin treatment desensitizes the position to mutation while Tunicamycin treatment causes sensitization. At position 63 we observe the opposite, with Tunicamycin desensitizing and Rapamycin sensitizing the position.

### Principal Component Analysis of Deep Mutational Scanning Data Across Chemical Perturbations

To reveal whether common responses were driving the differences between treatments, we performed principal component analysis on the difference fitness data (**Figure 3**). We focused on the first three principal components, which collectively explain 60 percent of the variance. Projecting the treatments onto the first two principal components shows that the sensitizing (Caffeine, Cobalt, DTT, HU, p-FP) and desensitizing treatments (Cerulenin, Menadione, MG132, Nickel, Tunicamycin) form independent clusters. Treatment with AmpB, Rapamycin, or Tamoxifen appear as outliers in this space.

**Figure 3:**
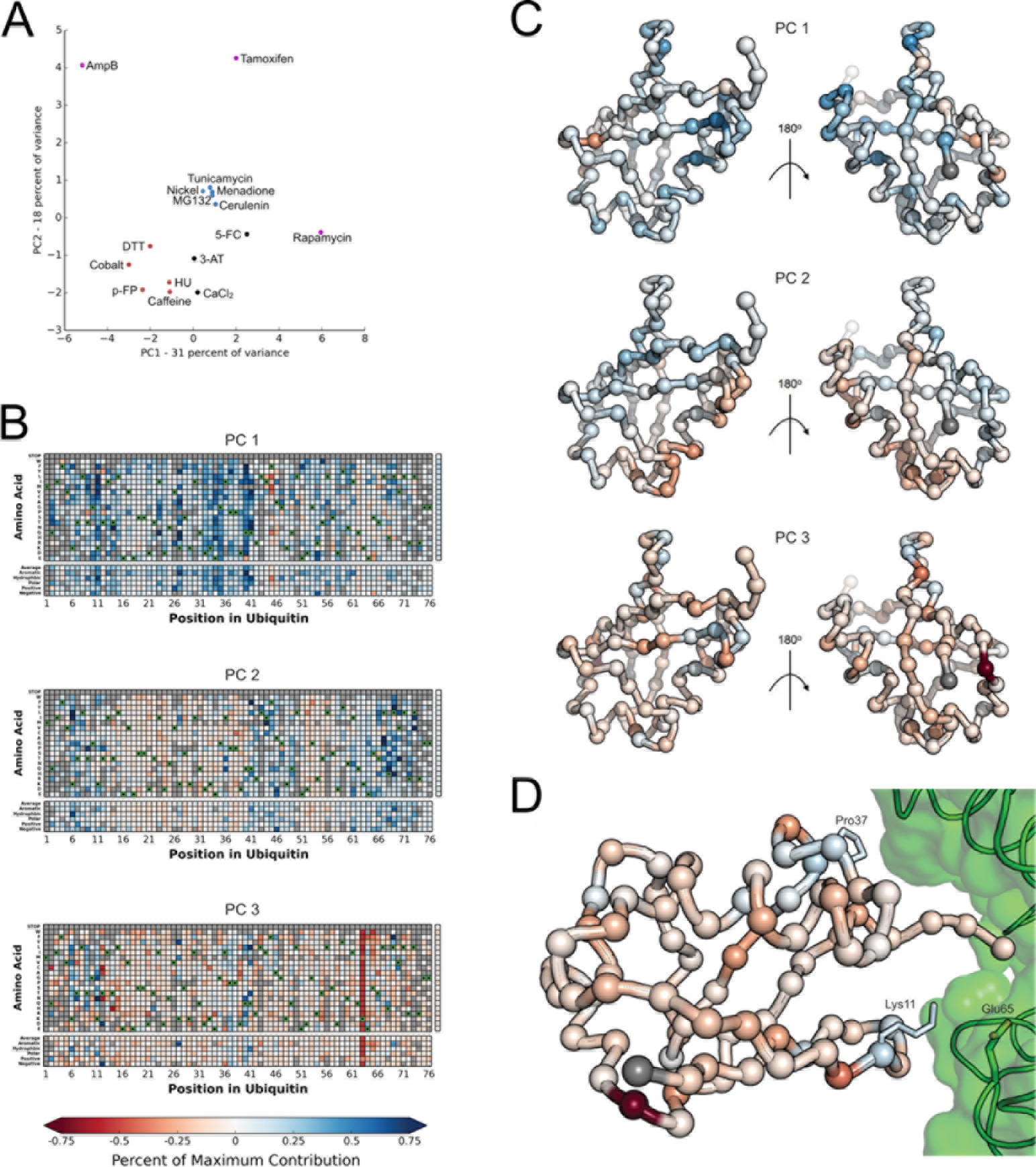
Principal component analysis reveals specific signals related to K63 incorporation: (A) The first two principal components reveal a sensitizing cluster plotted in red and an alleviating cluster shown in blue. Three strong outliers are shown in Purple. Treatment with 3-AT, 5-FC or CaCl_2_ appear between clusters with positive values for PC1 and negative values for PC2. (B) The contribution of each mutation to each principal component was visualized as a heat map. The percentage of the maximum contribution to that principal component is represented from 75% (Blue) to −75% (Red). (i) PC1 is related to the general sensitivity of Ub mutants to the perturbation. Negative values in the PC correspond to greater sensitization. Strikingly, this general sensitization is coupled to robustness to some mutations at the core residue Phe45. (ii) PC2 differentiates “Sensitive Face” residues (positive contributions) from “Tolerant Face” residues (Negative Contributions). (iii) PC3 reveals mutations that are correlated with sensitization of Lys63. (C) The average contribution of each mutation at each position was plotted from 75% (Blue) to −75% (Red) on the Ub monomer structure for PC1 (A), PC2 (B), and PC3 (C). (D) Rosetta ΔΔG calculations revealed that mutations that strongly destabilize the donor Ub pose on the MMS/Ubc13 (2GMI) heterodimer are localized to Lys11 and Pro37 (shown in sticks). In the case of Lys11, all mutations other than that to Arg destabilize the interface suggesting a salt-bridge between Lys11 and Glu65 of Ubc13 (shown in green sticks) Ubiquitin residues are colored by the contribution to PC3 as in panel C iii.

To examine the mechanistic basis for these clusters, we mapped the contribution of each mutation to each of the first three principal components (PCs). PC1 appears as mild positive contributions for most mutations, with the strongest positive signals appearing at residues 11, 27, 40 and 41. Interestingly, the strongest negative contributions appear at Phe45, a large core residue. PC2 separates the previously described tolerant face of Ub from the sensitive face, which includes the “hydrophobic patch”. Positions close to the hydrophobic patch residues have a positive contribution to PC2, while positions on the opposite face have negative contributions.

PC3 is dominated by the response to mutation at Lys63, a key poly-Ub linkage site. Lys63 linked poly-Ub is an important regulator of the DNA damage response in yeast and is also involved in efficient endocytosis and cargo sorting to the vacuole in yeast (Erpapazoglou et al., 2014). To investigate PC3, we used Rosetta to calculate the ddG of interaction for various complexes involved in Lys63 linked poly-Ub, including the closed and open forms of Lys63 linked di-Ub and the donor and acceptor ubiquitin poses on the MMS/Ubc13 heterodimer, the E2 that catalyzes Lys63 linked poly-Ub in yeast. We binned the calculated ddG values in stabilizing (ddG < −0.75) and destabilizing (ddG >= 0.75). We then calculated the contributions of each mutation to PC3 and compared the two distributions. Positions that destabilize this interface (mutations at Lys11 and Pro37) are enriched in positive contributions to the PC. This argues that conditions that require building K63-linked poly-Ub chains (those where position 63 is sensitized) have an increased relative fitness at mutations that destabilize the donor ubiquitin pose. Interestingly, when Lys11 is mutated to Arg, there is a strong negative contribution to PC3. This is the only mutation at position 11 that is predicted to stabilize the interface, likely due to Lys11 participating in a salt-bridge with Glu65 of Ubc13.

### Deep mutational scanning in different chemical environments reveals constraints on most residues

To continue our efforts in explaining the evolutionary constraints that lead to the exquisite conservation of Ub, we selected the condition with the minimum of average fitness at each position as the “minimum average fitness”. We then binned the minimum average fitness for each position into sensitive (≤−0.35), intermediate (−0.35 to −0.075) and tolerant (≥−0.075) (**Figure 4**). Previously we showed sensitization at all but twelve positions in Ub (Mavor et al., 2016). By expanding the number of perturbations, we now show all but two positions, Ser57 and Gln62, are sensitized in at least one condition.

**Figure 4:**
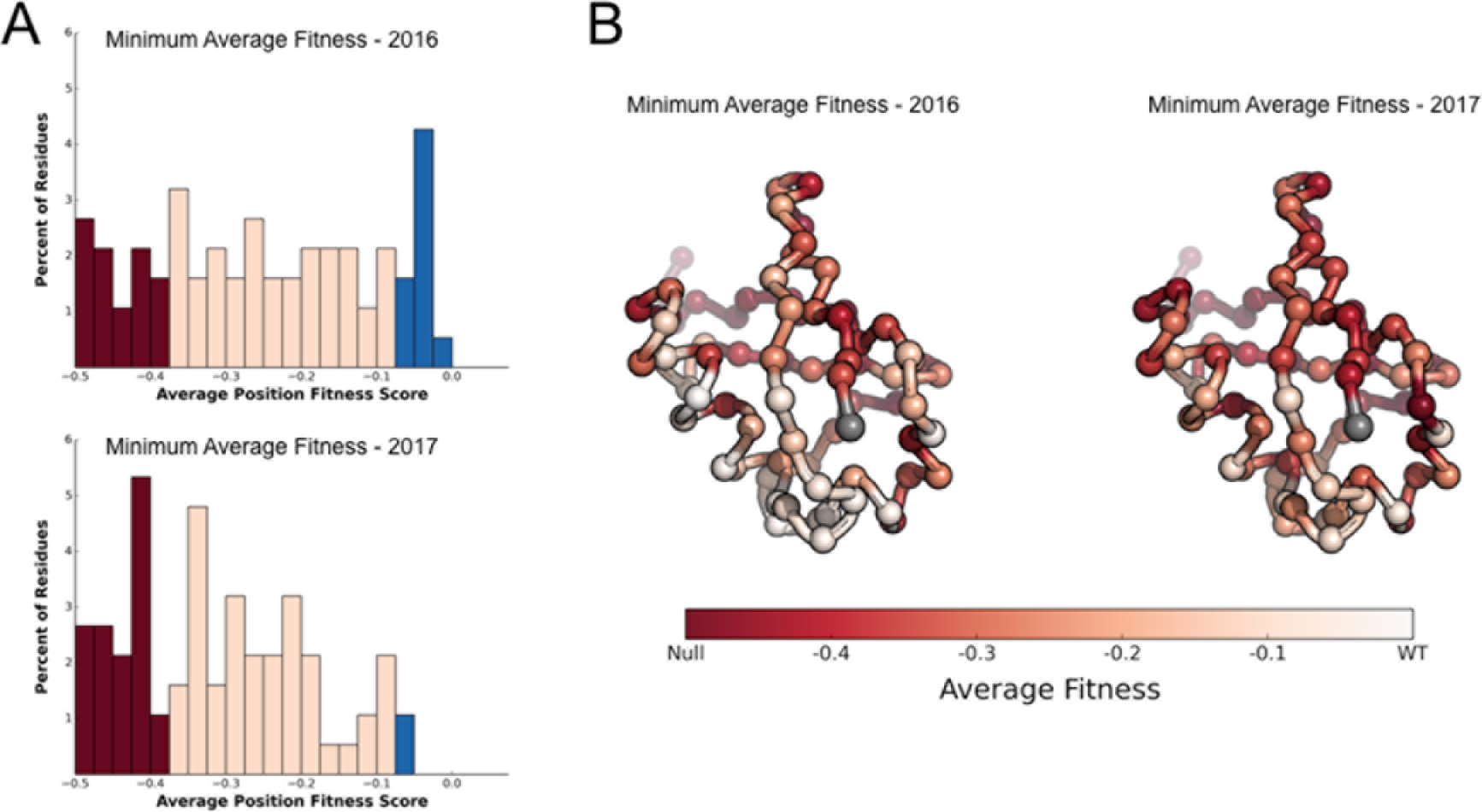
New perturbations reveal constraints on all but two Ub positions: (A) Positions were binned into tolerant (>=−0.075 − Blue), intermediate (<−0.075 to > −0.35 - Pink) and sensitive (<= −0.35 - Red) and the distributions plotted for each perturbation. Calculating the minimum average fitness reveals how the new perturbations reveal additional constraints on the Ub fitness landscape. (B) Average position fitness scores mapped onto ubiquitin. (i) Minimum average fitness score in DMSO, Caffeine, DTT, HU, and MG132 (ii) Minimum average fitness score in all conditions. C-alpha atoms are shown in spheres and the residues are colored according to average fitness. Met1 is colored grey.

## Discussion

No single perturbation can replicate the diverse pressures that natively constrain protein evolution. We can now rationalize many of the constraints that have led to the extreme sequence conservation of Ub by examining the fitness landscape under a large variety of conditions, including redox stress, osmotic stress, protein folding stress, DNA damage, ER stress, and anti-fungals. Notable exceptions are residues Ser57 and Gln62, which are not sensitive to mutation under any condition yet tested.

Of the newly revealed sensitivities, perhaps the most exciting is the sensitization of Lys63, captured by PC3, in eight conditions. Lys63-linked poly-Ub participates in the response to DNA damage, where Lys63-linked poly-Ub is catalyzed on PCNA to induce error-free postreplication repair (Zhang et al., 2011), and in endocytosis, where efficient endocytosis in cargo sorting to the vacuole requires Lys63-linked poly-Ub chains (Erpapazoglou et al., 2014).

In contrast, we previously observed an increase in mutational robustness at Lys63 in DTT treatment, a reducing agent that interferes with ER protein folding. Interestingly, we also observe increased robustness under Tunicamycin treatment, a compound that interferes with ER protein folding via a distinct mechanism (Chawla et al., 2011). This result suggests an epistatic interaction between Lys63 signalling and the unfolded protein response, which may complement the suggested role of Lys11 under high (30mM) DTT treatment (Xu et al., 2009). The Lys11Arg mutant is specifically sensitized in Tunicamycin suggesting that the origin of this effect may be structural, rather than due to a requirement for Lys11-linked poly-Ub.

In addition to the increased robustness at Lys63, Tunicamycin treatment leads to a unique increase in mutational robustness at several other positions, including Lys6, Lys11, and Lys33. These results address a major challenge in Ub biology: understanding the biological role of distinct poly-Ub species. While the mutational tolerance pattern at Lys6 and Lys11 appear to be due to disrupting a salt-bridge, the increased robustness at Lys33 suggests a connection between Tunicamycin and Lys33 linked poly-Ub. We observed, further, but less conclusive, Lysine-specific effects for Lys27, Lys29, and Lys33 under treatment with AmpB, Cobalt, or Nickel.

Finally, these experiments continue to highlight the success of project-based courses. Over the course of 6-8 weeks, first year graduate students in UCSF’s CCB and iPQB programs generated and analyzed these data using their own computational pipelines. We believe that the Ub yeast library is an ideal system for such project-based courses due to the wide range of stress responses mediated by the system and the vastness of chemical space. It is our hope that other graduate programs can offer similar project based classes in the future and our regents are available to further that goal.

## Methods

### Additional material is available

PUBS website (www.fraserlab.com/pubs)

GitHub (https://github.com/fraser-lab/PUBS).

Raw Sequencing reads are made available via SRA (SRA Accession Number:SRP070953)

### Updated Methods from Mavor et al 2016

For each compound, we determined the the chemical concentrations that inhibited SUB328 (WT Ub) growth by 25% (3-Amino-1,2,4-triazole - 50 mM, 5-fluorocytosine - 1.25 ug/mL, Amphotericin B - 400 nM, CaCl_2_- 500mM, Cerulenin - 4.5 uM, Cobalt Acetate - 600 uM, Menadione- 500 uM, Nickel Chloride - 400 uM, p-Fluorophenylalanine - 800 ug/mL, Rapamycin - 200 nM, Tamoxifen - 25 uM, and Tunicamycin- 1 mg/mL). Other growth, sequencing, and data processing methods are unchanged.

### Principal Component Analysis

PCA was performed using scikit-learn (version 0.18.1) in Python with the following parameters:

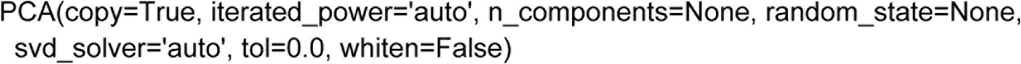

For each compound, the difference in fitness between DMSO and perturbation was calculated. PCA was performed on these 16 vectors. In the case of a missing observation for any mutant and positions, that mutant and position was excluded from the analysis.

### ROSETTA ddG predictions

Interface ddG predictions were generated using the Rosetta macromolecular modeling suite, which is freely available for academic use. The git version used was 12e38402d9. For each amino acid position in the MMS/Ubc13 heterodimer, the interface ddG protocol was run as follows: 1) Minimize (with constraints to the starting coordinates) the starting wildtype structure (PDB ID: 2GMI). 2) Generate an ensemble of 50 conformational states using Rosetta’s backrub application (10,000 trials, temperature 1.2), using residues in an 8Å radius of the specified amino acid position as backrub pivot residues. 3) Repack, or repack and mutate the side chains of the specified amino acid and the pivot residues from step 2. 4) Minimize (with constraints) the wild type and mutant structures generated in step 3. 5) For each structure *i* (of 50), we calculate the ddG score as follows:

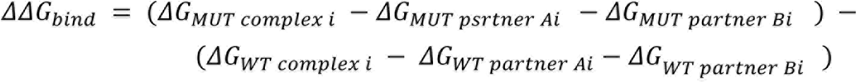

We then average all 50 *ΔΔG_bind_* scores to obtain the final predicted value.

## Acknowledgements

We acknowledge: administrative support from Rebecca Brown, Julia Molla, and Nicole Flowers; technical support from Jennifer Mann and Manny De Vera; gifts from David Botstein, and Illumina; and helpful discussions with Nevan Krogan and Ron Vale. The Project Lab component of this work is specifically supported by an NIBIB T32 Training Grant, ‘Integrative Program in Complex Biological Systems’ (T32-EB009383). UCSF iPQB and CCB Graduate programs are supported by US National Institutes of Health (NIH) grants EB009383, GM067547, GM064337, and GM008284, HHMI/NIBIB (56005676), UCSF School of Medicine, UCSF School of Pharmacy, UCSF Graduate Division, UCSF Chancellor’s Office, and Discovery Funds. WC, EMG, LRK, LSM, PS, TLN, NJR and FSU are supported by NSF Graduate Research Fellowships. DNB is supported by NIH GM112844. TK is supported by NIH R01 GM117189, R01 GM110089 and NSF MCB-1615990. HDM is supported by NIH R01 GM071801, R01 AI100272, R01 AI120464, R56 AI126726 and the Chan Zuckerberg Biohub. MK is supported by an NIH Director’s New Innovator Award (NIH/NIGMS DP2 GM119139), an Allen Distinguished Investigator Award (Paul G. Allen Family Foundation), a Stand Up to Cancer Innovative Research Grant, NIH K99/R00 CA181494, the Tau Center Without Walls (NIH/NINDS U54 NS100717), the Chan Zuckerberg Biohub and the Paul F. Glenn Center for Aging Research. JSF is a Searle Scholar, Pew Scholar, and Packard Fellow, and is supported by NIH 0D009180.

